# Inducing lucid dreaming with multisensory stimulation: a preregistered multi-center study

**DOI:** 10.1101/2024.06.21.600133

**Authors:** Claudia Picard-Deland, Mahdad Jafarzadeh Esfahani, Leila Salvesen, Tobi Matzek, Ema Demšar, Tinke van Buijtene, Victoria Libucha, Cosimo Cecconi, Bianca Pedreschi, Paul Zerr, Nico Adelhöfer, Emma Peters, Sarah F. Schoch, Giulio Bernardi, Michelle Carr, Martin Dresler

## Abstract

Lucid dreaming occurs when one is aware of dreaming while asleep. While lucid dreaming can occur spontaneously, it remains rare. This multi-center study (Netherlands, Italy, Canada) tested a combined induction approach integrating senses-initiated lucid dreaming (SSILD) with targeted lucidity reactivation in a large sample of participants with varying lucid dreaming experience.

Sixty participants (33 F; 26.5 ± 6.2 years old) completed two morning naps in the sleep laboratory. Participants received pre-sleep SSILD training paired with visual, auditory, and tactile cues that were then reintroduced in REM sleep, with stimulation and sham conditions counterbalanced. Lucidity and cue perception were verified using intentional eye movements from within the dream (signal-verified lucid dream, SVLD).

SVLDs occurred in 31 participants (51.7%) across both naps. Subjective lucidity occurred in 63 naps (52.5%), including 40 SVLDs (33.3%) with no difference between stimulation (38.3%) and sham (28.3%) conditions. However, SVLD duration and number of predefined eye movements were higher with sensory cueing. Cues were perceived within sleep in most cued REM periods (71.1%), sometimes acting as lucidity signals (39.1% of stim SVLDs) and occasionally disrupting sleep (23.7% of cued periods).

Overall, our combined cognitive-sensory induction method produced relatively high lucid dreaming rates, even in participants who rarely experience them. While cues were sometimes perceived as lucidity signals and may help prolong lucid episodes, our results do not show clear benefits of REM cueing over SSILD training alone. Further refinement of these methods may support clinical applications and deepen insight into consciousness and sensory processing during sleep.

## 1. Introduction

Lucid dreaming (LD), the state of becoming consciously aware of being in a dream, is a captivating, yet relatively rare phenomenon. While approximately half of the population has experienced spontaneous LD at least once in their lifetime, such occurrences remain quite infrequent (Saunders et al., 2016). Dream lucidity offers a distinctive opportunity for investigating the neuroscience of dreams by enabling the use of measurable lucidity verification techniques, such as predefined eye signals, breathing patterns, or facial muscle contractions (Baird et al., 2022; Holzinger et al., 2006; Konkoly et al., 2021; LaBerge et al., 1981). This objective signaling within LD offers improved experimental control over dream content and duration, enables external real-time monitoring, and even allows two-way communication with dreamers (Baird et al., 2019; Dresler et al., 2015; Konkoly et al., 2021; Siclari et al., 2017). Increasing levels of insight and control within dreams also hold promise for various clinical applications, including treatment of nightmare disorders (de Macêdo et al., 2019; Morgenthaler et al., 2018; Sandell et al., 2024), nightmare-related symptoms in post-traumatic stress disorder (PTSD)(Holzinger et al., 2020; Yount et al., 2024); narcolepsy (Rak et al., 2015) as well as for personal, recreational, and creative endeavors (Gott et al., 2020; Zink & Pietrowsky, 2013). A current challenge in dream research involves developing reliable methods to induce LD in everyday settings, clinical environments, and in the laboratory (Adventure-Heart, 2020; Mota-Rolim et al., 2019).

Previous efforts in LD induction encompass a wide range of approaches, including cognitive training (Adventure-Heart, 2020; Appel et al., 2020; Baird et al., 2019; Erlacher & Stumbrys, 2020; Schredl et al., 2020), external sensory stimulation during rapid eye movement (REM) sleep (Erlacher et al., 2020; Erlacher & Stumbrys, 2020; Paul et al., 2014), pharmacological interventions (Kern et al., 2017; LaBerge et al., 2018), brain stimulation (Blanchette-Carrière et al., 2020; Voss et al., 2014) and combinations of different methods (Adventure-Heart, 2020; Carr et al., 2023; Saunders et al., 2017; Schmid & Erlacher, 2020). A comprehensive review of induction techniques can be found elsewhere (Tan & Fan, 2023). While most approaches have shown little to moderate success rates at inducing objectively verified lucidity, targeted lucidity reactivation (TLR) has demonstrated the highest success rate in the laboratory to date (Carr et al., 2023). This induction method involves associating a pre-sleep lucidity training with sensory cues (visual and auditory) and then replaying the same cues during REM sleep. This TLR procedure elicited LD in 50% of participants, with objective lucidity verification using a predefined eye movement pattern to signal lucid episodes during the dream. Another promising cognitive technique is the senses-initiated lucid dreaming (SSILD) method, which involves cycling through the different senses while falling asleep (Zhang, 2013). This technique requires only a brief and easily implementable pre-sleep cognitive exercise, and demonstrated promising results in a home study (16.9% success rate across one week) (Adventure-Heart, 2020).

However, validation of this technique with physiological recordings is still necessary (Tan & Fan, 2023). The current body of research on LD induction techniques presents significant limitations, including small sample sizes, reliance on self-reported questionnaires in the absence of physiological data, and a focus on individuals who already experience LDs frequently (≥ 1 lucid dream episode per month) (Snyder & Gackenbach, 1988), restricting generalizability.

To address these limitations, we designed a study to validate a novel combination of LD induction techniques in a large, heterogeneous sample of participants with varying prior LD experience. Our induction method combines SSILD-based pre-sleep cognitive training with multimodal (visual, auditory, and tactile) TLR in subsequent REM sleep periods using commercially available wearable devices, a combination that has already shown promising results in case studies (Campillo-Ferrer et al., 2025). We compared the effects of SSILD training with and without additional REM cueing during morning naps in the laboratory using a within-subject design including N=60 participants, expecting that cueing during REM sleep would increase lucidity, overall dream awareness, and control.

## 2. Methods

This multicenter study involved data collection across three sleep laboratories in the Netherlands (NL), Italy (IT), and Canada (CA). The experimental protocol was preregistered in OSF (Esfahani, Carr, et al., 2022) prior to data collection in all participating centers. To obtain a pooled sample of N=60, each center recruited 20 participants who completed two morning naps with at least one REM period each, following a within-subject design (stimulation and sham conditions counterbalanced).

### 2.1 Participants

Participants were recruited through flyers, word of mouth, and online recruitment strategies specific to each research center. Inclusion criteria included being in general good health, aged between 18 and 55, having a regular sleep-wake pattern, having at least one prior LD experience, and a dream recall frequency of at least three times per week. Exclusion criteria included a presenting history of neurological, psychiatric, or neurodegenerative disorders, prior brain surgery, epilepsy diagnosis, pregnancy, and the use of sleep-altering medication. Participants were further screened for depression (score ≥ 20 from the Beck Depression Inventory II (BDI-II) (Beck et al., 1996)), anxiety (score > 15 from the Beck Anxiety Inventory (BAI) (Beck et al., 1988)), prodromal symptoms (score ≥ 9 from the Prodromal Questionnaire (PQ-16) (Ising et al., 2012)), sleep quality (score > 7 from the Pittsburgh Sleep Quality Index (PSQI) (Buysse et al., 1989)), and chronotype (sleep time before 23:00 or Rise time before 07:00 from the Morningness Eveningness Questionnaire (MEQ) (Horne & Ostberg, 1976). The compensation for the entire study was 70 EUR in European centers and 120 CAD in Canada. This study was approved by the institutional research ethics committee at each participating site. Written informed consent was obtained from all participants prior to their participation in the study.

A total of 131 participants were recruited (N=35 at the NL site; N=57 at the IT site; N=39 at the CA site) – of these, 24 were excluded due to above-threshold scores on baseline questionnaires, 33 did not have REM sleep in at least one nap, 7 dropped out after the intake session, and 7 were excluded due to technical difficulties (signal loss, high noise levels, or external disruption) (see Supplementary Figure S1 for more details). Recruitment continued until a final sample of 60 valid participants was obtained (N=20 from each site, age = 26.50 ± 6.15 yrs old; 33 F, 27 M), upon which all further analyses are based.

### 2.2 Study procedure

The overall study timeline and protocol are described in Figure 1. After an online or short phone screening process, eligible participants were invited to the intake session (in person at the NL and IT sites, online at the CA site), during which they received detailed information about the study and provided informed written consent. They then completed questionnaires prior to coming twice to the laboratory for two morning nap sessions, held approximately 1 and 2 weeks after the intake session, respectively. They were asked to keep a home dream diary starting 1 week before the first nap up until the second nap (a total of ∼2 weeks). Both nap sessions involved a pre-sleep SSILD training associated with a set of sensory cues. One of the morning naps also included cueing during subsequent REM sleep periods (stim), while the other did not (sham), with the order counterbalanced across participants. They were asked to signal lucidity and cue perception in real-time from within their dreams using a predefined eye movement pattern (left-right-left-right, LRLR) and to report any subjective experiences upon REM awakenings. To assess the short-term carryover effects of our induction technique on dreams at home, an optional survey was sent 2 weeks after the last experimental session, asking about any perceived changes in dreaming or waking life following the study.

**Figure 1.**
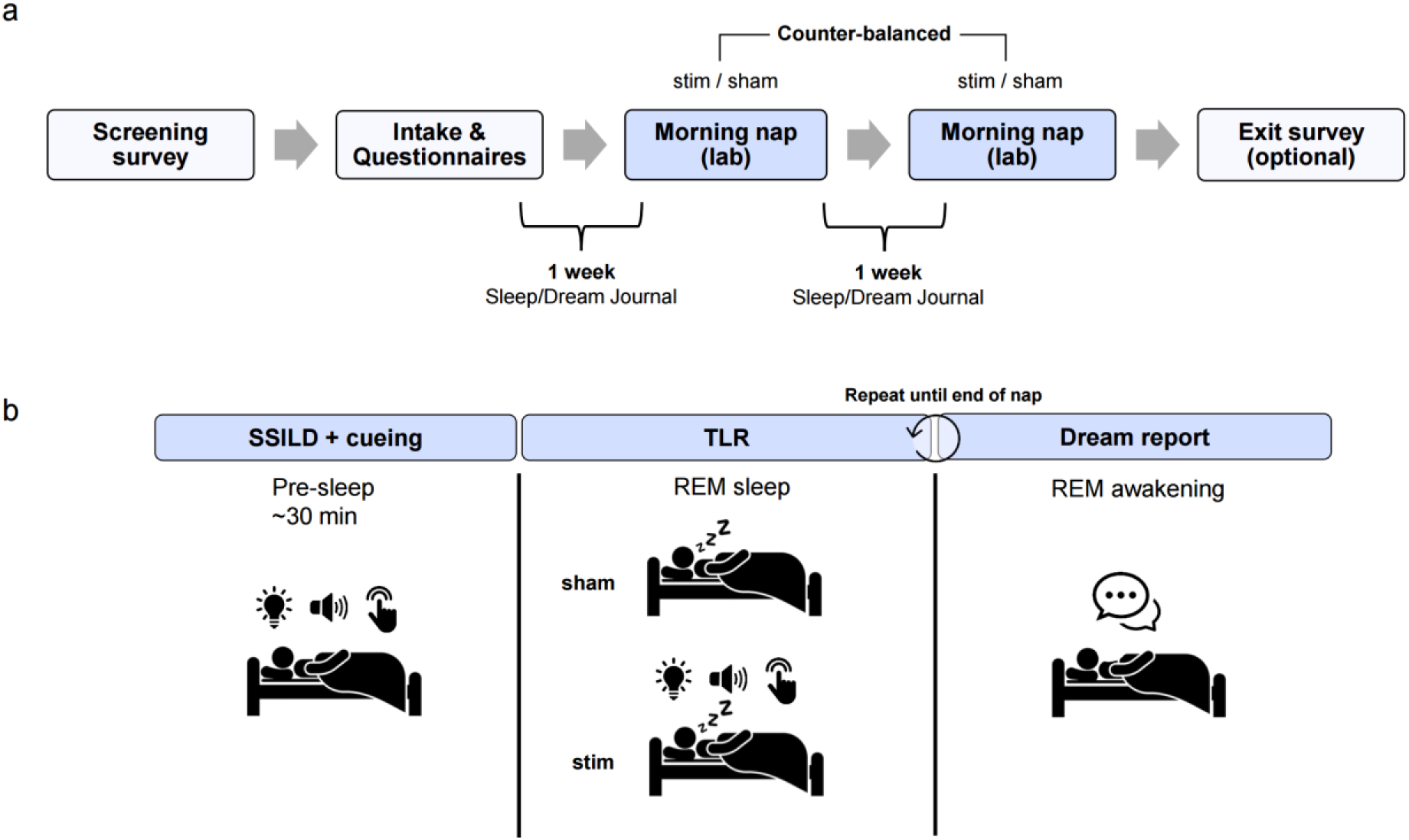
Overview of the study timeline and experimental protocol. (a) Following screening and intake, participants completed questionnaires, maintained a home dream diary for two weeks, and took part in two counterbalanced morning nap sessions (stim and sham conditions). An optional exit survey assessed perceived short-term effects after study completion. (b) Each nap included pre-sleep SSILD training with sensory cues. In one nap, cues were presented again during REM sleep (targeted lucidity reactivation; stim condition), whereas no cueing occurred in the other nap (sham condition). Lucidity onset and cue perception were signaled during sleep using predefined eye movements. Participants were awakened after each REM period, and dream reports were collected using a semi-structured interview and DLQ scores.

#### 2.2.1 At-home sleep and dream diaries

Participants were required to document their sleep-wake schedule and subjective assessment of sleep quality for the entire duration of the study. Dream reports and a Dream Lucidity Questionnaire (DLQ) (Stumbrys et al., 2013) were also collected daily. Additional questionnaires collected information on retrospective sleep and dreams (Mannheim Dream Questionnaire - MADRE) (Schredl et al., 2014) and waking cognition (Vividness of Visual Imagery Questionnaire - VVIQ) (Marks, 1973) (not analysed here). Home diaries and questionnaires were accessed online via Castor EDC (NL and IT sites) or REDCap (CA site) web platforms.

#### 2.2.2 Experimental nap protocol

Participants arrived at the lab at 07:00 a.m. at the NL and CA sites and between 05:00 - 08:00 a.m. in IT, depending on participants’ usual sleep-wake schedule and laboratory availability. They were requested to refrain from consuming any alcoholic or caffeinated beverages in the evening and morning before the nap sessions. Both naps followed the same pre-sleep lucidity training (see next section), after which participants were allowed to sleep for up to 2.5 hours. Participants were told that both naps might or might not be cued to remain blind to the experimental condition. Continuous background white noise was played throughout the naps at ∼35 dB (level adjusted to participants’ comfort) to soften the impact of auditory cueing and standardize the auditory environment across laboratories.

#### 2.2.3 SSILD training

Prior to their nap, participants underwent a ∼30-min lucidity training based on the SSILD method (see Supplementary Material for script). A pre-recorded audio track (English in NL, Italian in IT, French or English in CA) guided participants through a series of SSILD cycles, during which they were instructed to lie in bed with their eyes closed and focus their attention towards each sensory modality at a time (vision, hearing, and bodily sensations), starting with 4 fast cycles (2-3 s on each modality), followed by 4 medium length cycles (20 s), and ending with 4 slower cycles (60 s). During the slower cycles, each sensory modality was associated with its corresponding cue, with intensity decreasing through each cycle. The visual cue consisted of 3 brief red-light flashes from the ZMax headband; the auditory cue consisted of a 3-tone melody played through loudspeakers (500-700-900 Hz); and the tactile cue consisted of 1 to 3 soft vibrations on the forehead from the ZMax headband (see Supplementary Table S1 for cueing protocol during training). A vocal prompt reminded the participant to practice a lucid mindset (as in Carr et al., 2023) each time they perceived a cue during the first 2 slow cycles; cues were then presented alone without prompts for the last 2 slow cycles, after which the participant could fall asleep normally (see Supplementary Figure S2 for a visual overview of the cognitive training). The cues were maintained throughout the entire training, even if the participant fell into light sleep; however, the cues were stopped if the participant fell into REM sleep.

#### 2.2.4 REM cueing protocol

In the stimulation condition, the sensory cues presented during the SSILD training were presented again in each subsequent REM period. Cueing started ∼20 seconds after detecting REM sleep, and cues were presented in the same order as during the SSILD training (i.e., visual, auditory, and vibration) approximately every 20 seconds.

Visual and vibration cueing started with the minimum intensity levels afforded by the ZMax headband, or with the minimum light perception threshold if higher; auditory cueing started at 40dB or at the minimum subjective threshold, if lower (see Supplementary Methods for the evaluation of individual stimulation levels). Cue intensities increased incrementally during REM sleep: by 5% for light, 5 dBA for sound, and from one to three repetitions for vibration. If signs of microarousal (e.g., elevated EMG or alpha activity) were detected, stimulation paused and resumed only after full dissipation of arousal signs, with cue intensities reset to the level preceding the disturbance. If the participant responded with intentional LRLR eye movements, intensities were held constant until awakening. All cues were reset to baseline levels for the next cueing period.

#### 2.2.5 Lucid dream instructions

Participants were asked to do an intentional predefined eye movement sequence (i.e., LRLR) in the following cases: (1) whenever they became aware of the fact they were dreaming, allowing for an objective marker of the “initiation” of the lucid episode; (2) each time a cue was perceived during sleep, providing a marker for stimulus perception within the dream (cue incorporation); (3) approximately every 30 seconds if no sensory cue was perceived, to estimate the duration of the lucid dream.

Participants were also informed of the high probability of dreaming about the sleep laboratory or experiencing false awakenings. In such cases, they were recommended to perform a reality check (e.g., breathe through a pinched nose or count fingers) whenever unsure of their state. In case they became lucid, they were advised to calmly observe or explore the dream scene and avoid intense actions (e.g., flying or free-falling) during the initial moments of lucidity to increase the likelihood of sustaining the lucid state.

#### 2.2.6 Dream interview

At the end of each REM period (upon spontaneous awakening or at the first signs of another sleep stage), participants were awakened to report any subjective experience recalled from the last moments before waking, including sensations, feelings, thoughts, or emotions. They were also asked whether they felt they were asleep during the experience, whether they were aware of their conscious state, and to provide additional details when possible (e.g., estimated duration, precise content, cue perception, presence of eye signaling, following the dialogic structure in Supplementary Figure S3). Participants then completed the DLQ based on the reported experience, if any. After completing the dream interview, participants were invited to go back to sleep, if possible, for the remainder of the 2.5-hour session. They first listened to a shorter version of the SSILD training (∼ 7-min) consisting of a few minutes of verbal uncued fast and slow cycles, followed by a single cued cycle without verbal prompts, before resuming sleep (see Supplementary Material for the script and Table S2 for the cueing protocol in the short training).

Dream reports were also collected at the end of the nap (after 2.5 hours had elapsed), upon a spontaneous post-LD awakening, upon observing an LRLR signal during non-REM sleep, or upon experimental interruptions such as bathroom breaks or technical adjustments. Upon final awakening, participants completed the LuCiD scale questionnaire (Voss et al., 2013) for each REM awakening from which a dream report was collected.

### 2.3 Signal acquisition and sleep scoring

As the study aimed to be a proof-of-principle that LD induction is possible also on a larger scale, such as clinical or home settings, electroencephalographic (EEG) measurements were collected using ZMax EEG headbands (Hypnodyne Corp., Sofia, Bulgaria), a validated sleep wearable device (Esfahani et al., 2023) equipped with two frontal EEG channels (F7-Fpz, F8-Fpz). Three additional electromyographic (EMG) channels recorded muscular activity from the chin area using site-specific software and systems. At the CA site only, an additional standard polysomnography (PSG) montage was applied (EEG: F3, F4, C3, C4, O1, O2; 2 horizontal EOG channels) (see Supplementary Methods for site-specific systems).

The open-source dream engineering toolbox Dreamento (Esfahani, Daraie, et al., 2022) was used for real-time EEG signal monitoring, recording, sensory stimulation, and offline data analysis. The ZMax signals were also recorded through the manufacturer’s proprietary software (as described in Esfahani et al.)(2022). A calibration period was conducted at the start of each recording to establish baseline measures and synchronize signals (see Supplementary Methods for details).

For the NL and IT sites, sleep stage scoring was performed using the ZMax signal with offline Dreamento by two researchers from the other sites (see Supplementary Methods for inter-rater agreements details); for the CA site, sleep scoring was performed by a sleep expert using the full PSG signal (see Esfahani et al., 2023 for a full validation of ZMax signals against PSG). Sleep stage scoring was used to validate that the awakenings, stim/sham periods, and lucid signals included in the analyses occurred in REM sleep. Sleep metrics were evaluated using the YASA Python toolbox (Vallat & Walker, 2021).

### 2.4 Lucid dream classification

A lucid dream was considered “subjective LD” if the participant reported becoming lucid and/or indicated being at least “moderately” aware of being in a dream (i.e., score -≥2 on a scale of 0-4 from the first question of the DLQ, “I was aware that I was dreaming”). A signal-verified lucid dream (SVLD) was confirmed in the presence of both objective and subjective measures of lucidity, i.e., when 1) a predefined eye movement (LRLR) was detected and 2) the subject reported becoming lucid (as defined above) and reported performing the signal.

Predefined eye movements were identified on ZMax recordings by three scorers for each nap (one scorer from each site: MJE, LS, CPD/TM). If the participant performed intentional eye movements different from the predefined LRLR signal (e.g., partial, such as LRL), it was accepted as a lucidity signal only if the participant reported attempting to perform the LRLR signal. Only signals that were identified by at least 2 out of 3 scorers were considered. Predefined eye movements that occurred within 5 seconds after a sensory cue were considered to be cue-related; otherwise, they were considered cue-unrelated.

The duration of SVLD was estimated in three ways: 1) from the first predefined eye signal to the subsequent awakening; 2) from the first to the last predefined eye signal within the same REM period (subsample of SVLDs with ≥2 predefined eye signals) and 3) by considering only intervals between consecutive eye signals separated by less than one minute (i.e., continuous SVLD duration, subsample of SVLDs with ≥2 predefined eye signals).

Average and max scores of DLQ (DLQ-item 1 awareness, DLQ-item 4 control, DLQ total score) and LuCiD scale (total score and subscales: insight, control, thought, realism, memory, dissociation, negative emotion, positive emotion)(Voss et al., 2013) were also used to assess the level of lucidity for each nap.

### 2.5 Qualitative analyses of dream reports

We evaluated the occurrence of other dream experiences often associated with LD, including incorporation of the laboratory in dreams (Picard-Deland et al., 2021), false awakenings (Buzzi, 2011), sleep misperception (Bastien et al., 2014), sleep paralysis (Denis, 2018) and out-of-body experiences (Alvarado, 1992)(see Supplementary Methods for full definitions).

Dream reports collected in the laboratory were scored for each of these categories, as well as for cue incorporation, i.e., whether the sensory cues were perceived during sleep, and if so, whether they were incorporated into the dream narrative. Dream scorings were first done by one judge per site, then verified by one judge across all sites for consistency.

### 2.6 Statistical and methodological evaluation

Statistical analyses and figures were performed using R. All main analyses were conducted using linear mixed models (LMM) or generalized linear mixed models (GLMM), with condition (stim, sham) and site (CA, IT, NL) as fixed effects and subject as a random intercept. LMMs were used for continuous outcomes and GLMMs with a binomial distribution used for binary outcomes and count data. Fixed effects were tested using Type II Wald chi-square tests. Post-hoc pairwise comparisons were conducted using estimated marginal means with Tukey correction for multiple comparisons. Between-subject comparisons on prior lucid dreaming experience were conducted using Mann-Whitney U tests and Kruskal-Wallis tests were used to compare eye signal response times as a function of the 3 different cue modalities. The LuCiD scale used in the CA site originally ranged from 1 to 5, while it ranged from 0 to 5 in other sites; the CA LuCiD scores were thus proportionally transformed to fit the 0 to 5 scale before statistical analyses. When multiple REM awakenings occurred within a nap, responses were averaged across awakenings, and nap-level averages (or max values when stated) were used for subsequent statistical models. All statistical tests were 2-tailed, at α = 0.05.

## 3. Results

### 3.1 Prevalence of lucid dreams

Overall, 44 of 60 (73.3%) participants experienced at least one subjective LD, and 31 out of 60 (51.7%) experienced at least one SVLD over the two naps. A total of 63 out of 120 (52.5%) naps contained at least one subjective LD: 30/60 (50.0%) in the stim condition, and 33/60 (55%) in the sham condition. Neither condition (χ²(1) = 0.34, p = .562) nor laboratory site (χ²(2) = 0.73, p = .693) predicted subjective LD prevalence *(naps with LD ∼ condition + site + (1|subject))*. At least one SVLD occurred in 40 out of 120 (33.3%) naps: 23/60 (38.3%) in the stim condition, and 17/60 (28.3%) in the sham condition. Neither condition (χ²(1) = 1.51, p = .219) nor laboratory site (χ²(2) = 1.85, p = .396) predicted SVLD prevalence *(naps with SVLD ∼ condition + site + (1|subject))* (Figure 2).

**Figure 2.**
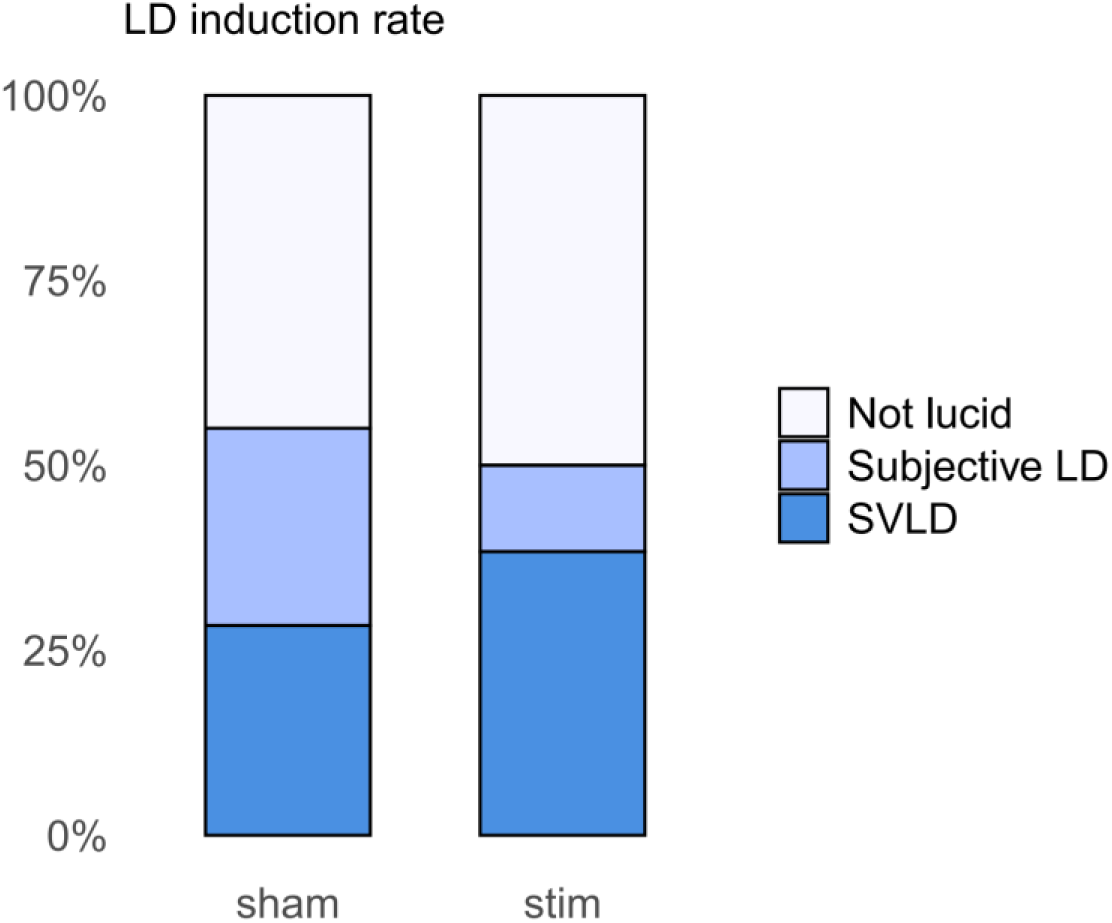
Prevalence of lucid dream experiences across the two conditions. Proportion of naps containing no lucid dreaming (Not lucid), at least one subjective lucid dream (Subjective LD), and at least one signal-verified lucid dream (SVLD) in the sham and stim conditions. Subjective LDs were based on self-reported ratings of lucidity, whereas SVLDs additionally required predefined eye signals during REM sleep. Percentages are calculated across all naps in each condition (60 naps per condition). No significant differences were observed between the stim and sham conditions for either subjective LDs or SVLDs.

In the stim condition, there were a total of 36 subjective LDs across 97 valid REM awakenings (37.1%; from 29 participants); in the sham condition, there were a total of 34 subjective LDs across 98 valid REM awakenings (34.7%; from 33 participants). Neither condition (χ²(1) = 0.13, p = .715) nor laboratory site (χ²(2) = 0.50, p = .779) predicted the proportion of LD trials per nap (*% LD trials per nap ∼ condition + site + (1|subject))*. When only considering SVLDs, 23 (23.71% of 97 REM awakenings; from 21 participants) occurred in the stim condition, compared with 17 (17.34% of 98 REM awakenings; from 17 participants) in the sham condition. Neither condition (χ²(1) = 1.35, p = .245) nor laboratory site (χ²(2) = 1.83, p = .400) predicted the proportion of SVLD trials per nap (*% SVLD trials per nap ∼ condition + site + (1|subject))*. Of note, 3 additional SVLDs in the stim condition occurred in REM periods when no cues were delivered (non-valid awakenings). These 3 SVLDs were included in the nap-level analyses as stimulation was presented during other REM periods in the same nap.

In addition to REM lucidity, there were also 13 SVLDs that occurred in NREM stages (6 in N1 and 7 in N2); 8 occurred in stim naps, and 5 occurred in sham naps. As only two of the N2 cases were confirmed with PSG scoring, these NREM-stage classifications should be interpreted with caution; these SVLDs were not included in the analyses. Finally, there were a total of 11 eye-signaled dreams in REM sleep without reported lucidity, including 9 in stim naps (6 within 5 seconds of a cue, including 1 who perceived the stimuli and remembered signalling, but did not get lucid, 1 without memory of doing the predefined eye signal, and 4 who felt like they were not asleep; and 3 occurring more than 5 seconds after a cue, including 1 who was unsure if they were dreaming, and 2 who perceived the stimuli and remembered signalling but did not get lucid), and 2 in sham naps, both without memory of doing the predefined eye signal.

#### 3.1.1 Level of lucidity

Linear mixed models were used to predict DLQ and LuCiD scores (*scores ∼ condition + site + (1|subject)*).

*DLQ - Awareness*. Neither condition nor laboratory site predicted *average awareness* score (χ²(1) = 0.16, p = .694; χ²(2) = 0.30, p = .859) or *max awareness* score (χ²(1) = 0.07, p = .789; χ²(2) = 0.04, p = .980).

*DLQ - Control*. Condition did not predict *average control* score (χ²(1) = 0.81, p = .368) or *max control* score (χ²(1) = 0.73, *p* = .394). Laboratory site predicted both *average* (χ²(2) = 6.71, p = .035) and *max control* scores (χ²(2) = 6.55, p = .038). Post-hoc comparisons for *average control* revealed lower scores for the IT site (M = 0.91) than the NL site (M = 1.80) (*t*(56.6) = −2.47, p = .044), while no other pairwise comparisons were significant. For *max control* score, marginal means suggested lower scores in the IT site (M = 1.17) compared to the CA site (M = 2.10) and NL (M = 2.12), though no pairwise comparison survived Tukey correction. *DLQ - Total score*. Neither condition nor laboratory site predicted *average total* score (χ²(1) = 0.34, p = .560; χ²(2) = 0.85, p = .655) or *maximum total* DLQ score (χ²(1) = 0.06, p = .811; χ²(2) = 1.69, p = .430) (Figure 3).

**Figure 3.**
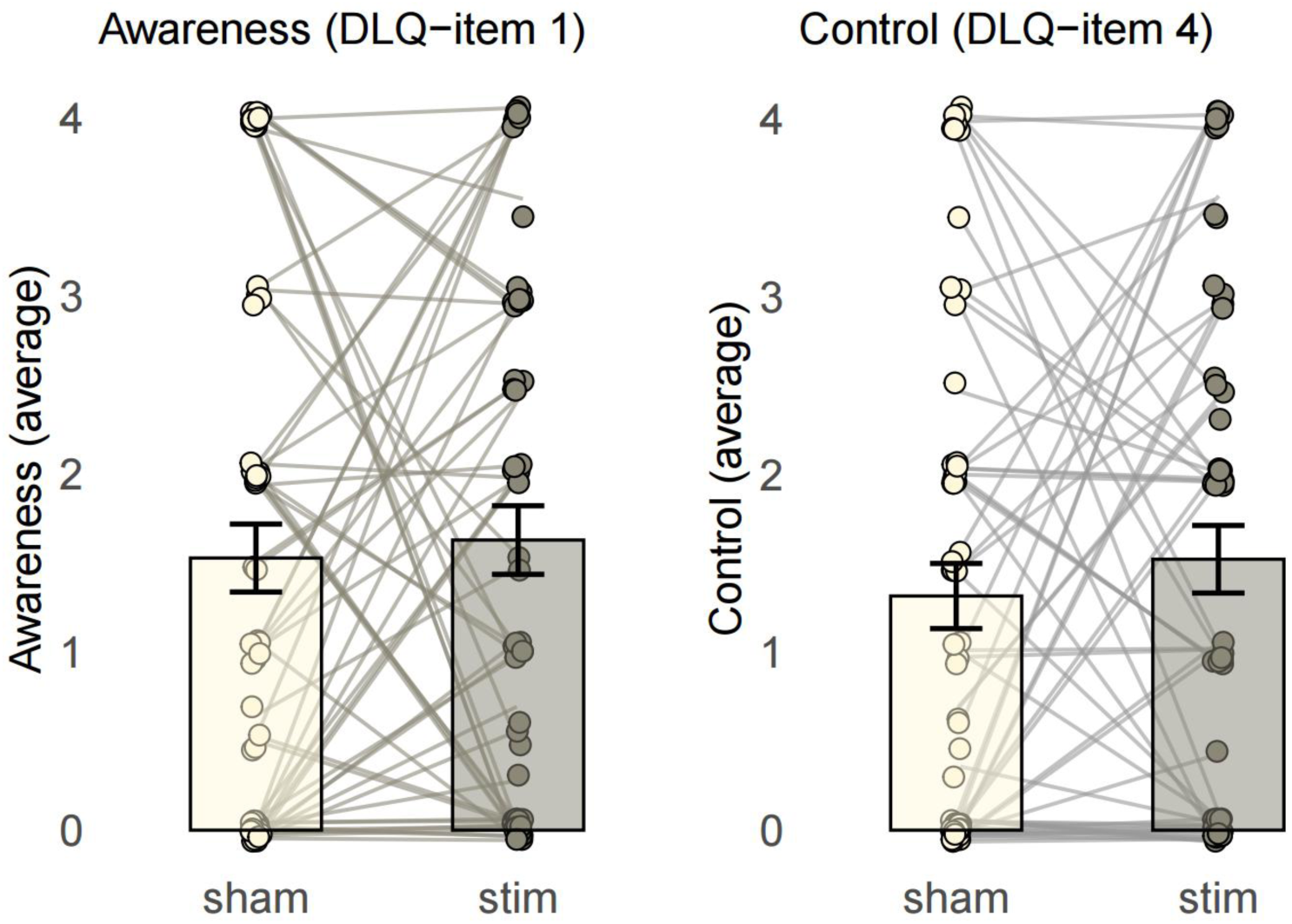
Dream Lucidity Questionnaire (DLQ) scores for Awareness (item 1) and Control (item 4) averaged across conditions. Within-subject differences in DLQ ratings are shown for the stim and sham conditions (mean ± SEM). Awareness was assessed with item 1 (“*I was aware that I was dreaming*”) and Control with item 4 (“*I deliberately chose one action instead of the other*”), scored on a 0-4 scale (0 = not at all; 1 = just a little; 2 = moderately; 3 = pretty much, 4 = very much). No significant differences were observed between stim and sham conditions for awareness, control, or total DLQ scores.

*LuCiD subscales.* All LuCiD subscales (insight, control, thought, realism, memory, dissociation, negative, positive), both average and maximum values, did not significantly differ by condition (all p > .060) or site (all p > .078), with only a marginal trend observed for higher *maximum control* scores in the stim condition compared to the sham condition (χ²(1) = 3.53, p = .060).

#### 3.1.2 Lucidity duration

Three measures of SVLD duration were examined using linear mixed models *(duration ∼ condition + site + (1|subject)).* First, duration from the *first LRLR to subsequent awakening* (either spontaneous or elicited at the end of the REM period), had an overall mean of 7.67 ± 7.64 min (median ± IQR: 5.85 ± 6.86 min), ranging from 7 s to 30.25 min; and did not significantly differ by condition (stim: 7.60 ± 7.86 min; sham: 7.78 ± 7.57 min; χ²(1) = 0.04, p = .836) or site (χ²(2) = 1.95, p = .377). Second, duration from the *first to last LRLR* (in cases when at least two LRLRs occurred) was longer in stim SVLDs (6.08 ± 6.98 min, ranging from 12 s to 29.72 min) than in sham SVLDs (3.24 ± 5.43 min, ranging from 15 s to 17.88 min; χ²(1) = 4.91, p = .027), with no significant effect of site (χ²(2) = 4.63, p = .099). Third, *continuous SVLD duration*, (considering the distance only between LRLRs occurring within 1 minute of each other) did not differ between conditions (stim: 76.84 s ± 53.99 s; sham: 47.50 s ± 25.39 s; χ²(1) = 2.31, p = .129) or site (χ²(2) = 0.13, p = .938) (Figure 4).

**Figure 4.**
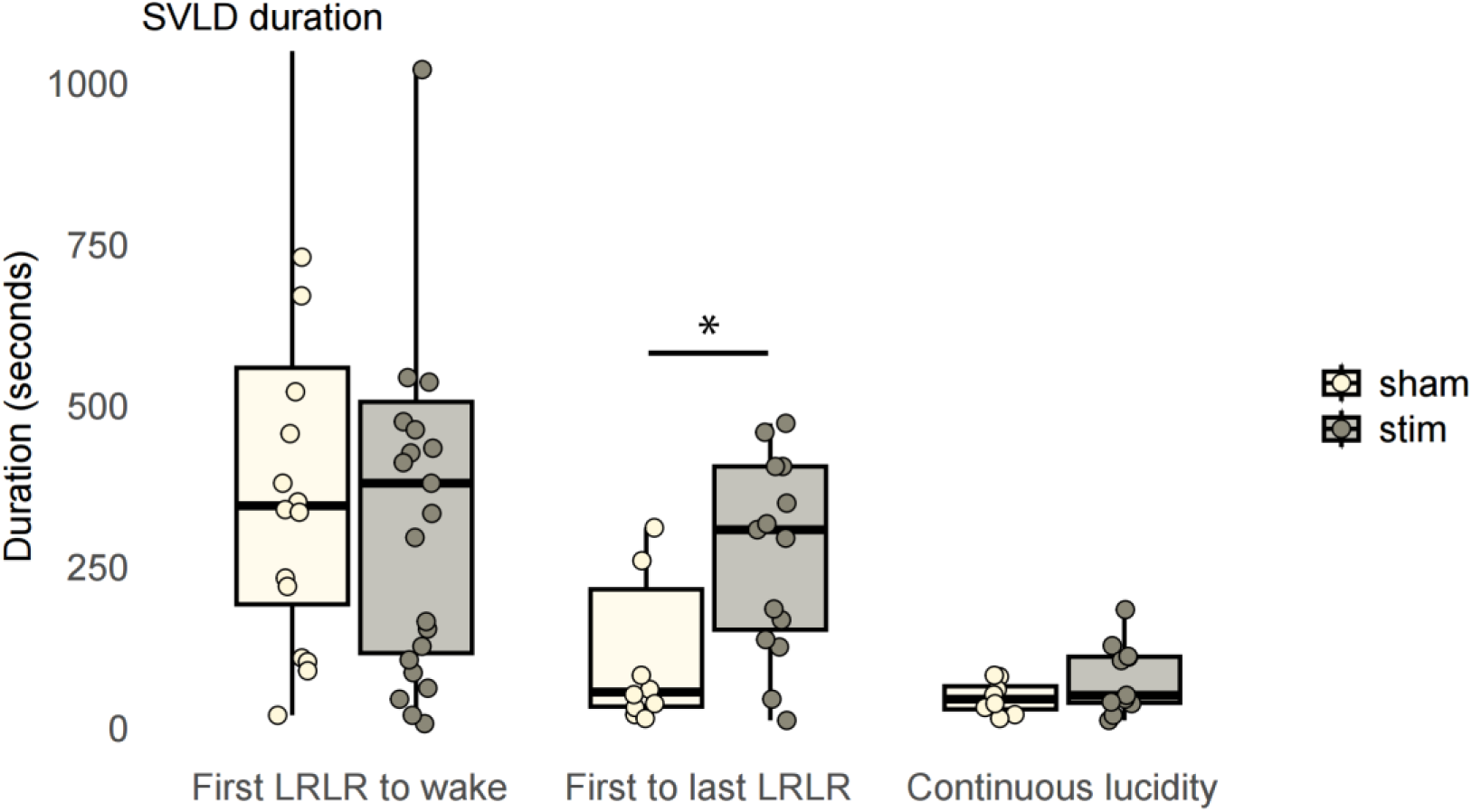
SVLD estimated duration (in seconds). The duration of SVLDs was estimated using three complementary approaches: 1) from the first predefined eye signal (LRLR) to the subsequent awakening; 2) from the first to the last LRLR within the same REM period (subsample of SVLDs with ≥2 predefined eye signals) and 3) a continuous estimate based only on LRLRs occurring within 1 minute of each other (subsample of SVLDs with ≥2 predefined eye signals). The interval between first and last LRLR was longer in the stim condition than in sham. *p<0.05

#### 3.1.3 Prior lucid experience

Participants who had at least one subjective LD in the lab across the two naps (with or without eye signals) had significantly higher prior LD frequency (MADRE score 3.70 ± 1.76 ≈ about 2-4 times a year) compared to those who didn’t (5.06 ± 0.85 ≈ about once a year) (Mann-Whitney U-Test W=516.5, p = .005). Participants who successfully signaled lucidity (SVLD) in the lab had marginally higher prior lucid dream frequency (3.74 ± 1.69 ≈ about 2-4 times a year) compared to those who didn’t (4.41 ± 1.62 ≈ about 2-4 times a year) (Mann-Whitney U-Test W=560.5, p = .094) (see Figure 5). Of note, 10 participants who reported only having lucid dreams once a year or less were able to achieve an SVLD in the laboratory.

**Figure 5.**
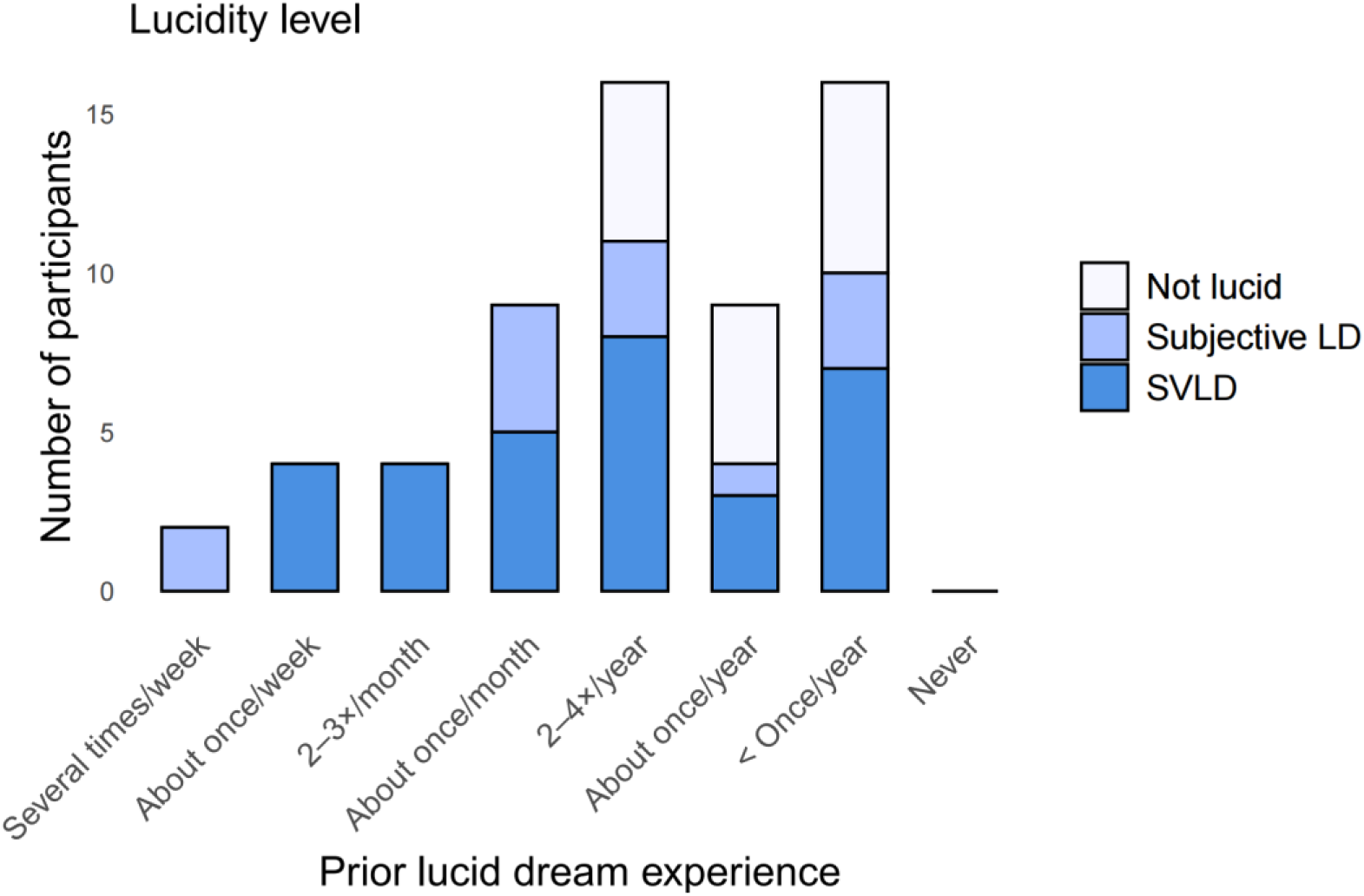
Lucidity level across prior lucid dream experience. Number of participants who had at least one subjective LD experience or at least one SVLD across the two naps, grouped by their prior lucid dreaming experience. Participants who reported at least one subjective LD in the laboratory (with or without eye-signaled lucidity) had higher prior lucid dream frequency than those who did not, whereas prior LD frequency was only marginally higher in participants who achieved SVLD. Notably, several participants with infrequent prior lucid dreaming were still able to achieve SVLD in the laboratory.

#### 3.1.4 Prevalence of LD-related dream experiences

The incorporation of the laboratory in dream narratives occurred in 67 naps (55.8%; 32 stim, 35 sham); false awakenings occurred in 56 naps (46.7%; 28 stim, 28 sham); sleep paralysis occurred in 8 naps (6.7%; 4 stim, 4 sham); and sleep misperception occurred in 37 naps (30.8%; 22 stim, 15 sham). The prevalence of these dream experiences did not differ by condition (all p > .154). Laboratory site predicted sleep misperception prevalence (χ²(2) = 6.11, p = .047), with estimated probabilities suggesting lower prevalence at the CA site (p = .140) compared to IT (p = .390) and NL (p = .370), though no pairwise comparison survived Tukey correction (all p > .051). Laboratory site did not predict any other dream experiences (all p >

.388).

#### 3.1.5 Predefined eye signals

Across all participants, a total of 154 predefined eye signals were identified, including 116 in the stim condition and 38 in the sham condition. In the stim condition, 73 were in response to cues (i.e., within 5 seconds after the cue: 27 to visual cues, 21 to auditory cues, and 25 to tactile cues) and 43 were cue-unrelated (i.e., not occurring within 5 seconds after any cue). Linear mixed models *(#LRLR ∼ condition + site + (1|subject))* were used to predict the number of eye signals. On average, the total number of predefined eye signals was higher in stim naps (1.93 ± 4.38) than in sham naps (0.63 ± 1.21) (χ²(1) = 5.04, p = .025), while site had no significant effect (χ²(2) = 0.45, p = .800). The number of cue-unrelated predefined eye signals did not differ between conditions (stim: 0.72 ± 1.85, sham 0.63 ± 1.21; χ²(1) = 0.09, p = .762) or site (χ²(2) = 0.24, p = .886). Within SVLDs, the total number of predefined eye signals was also higher in stim SVLDs (4.26 ± 5.82) than in sham SVLDs (2.12 ± 1.41) (χ²(1) = 8.23, p = .004). The response time to the sensory cues was of 2.18 ± 1.07 s on average (visual: 2.9 ± 0.12 s, auditory: 1.99 ± 0.58 s, tactile: 1.50 ± 0.64 s), which differed between modalities (χ²(2) = 12.92, p = .002), being shorter after tactile than after visual cues (p = .003, see Supplementary Figure S4 for a sample LRLR signal as seen on EEG).

#### 3.1.6 Perception of sensory cues

In the stim condition, cues were perceived to some extent by the sleeping participant in 69 out of 97 (71.1%) REM stim periods: in 9 (9.3%) stim periods the participants perceived the cues in their sleep, in absence of an ongoing dream (6 visual (V), 5 auditory (A), 4 tactile (T)); in 12 (12.4%) stim periods the participants perceived the cues (6 V, 9 A, 6 T) but felt awake (sleep misperception); and in 49 (50.5%) stim periods the cues were perceived during an ongoing dream. Among these, there were 30 (30.9% of all REM awakening) instances where at least some of the cues were incorporated and integrated into the dream narrative (19 V, 11 A, 17 T; see examples in Figure 6), in contrast to 22 (22.7%) cases where some cues were perceived as such within the dream (i.e., recognised as external stimuli without being integrated or transformed into the scenario). There were an additional 2 (2.1%) stim periods (including 1V, 1A and 2T) in which participants reported that their dream “paused” during the cues and the content resumed afterwards. It is of note that multiple types of incorporation could be present within one stim period, for example, a participant could report a single dream in which some cues were integrated into the dream narrative, while others were perceived as external stimuli.

**Figure 6.**
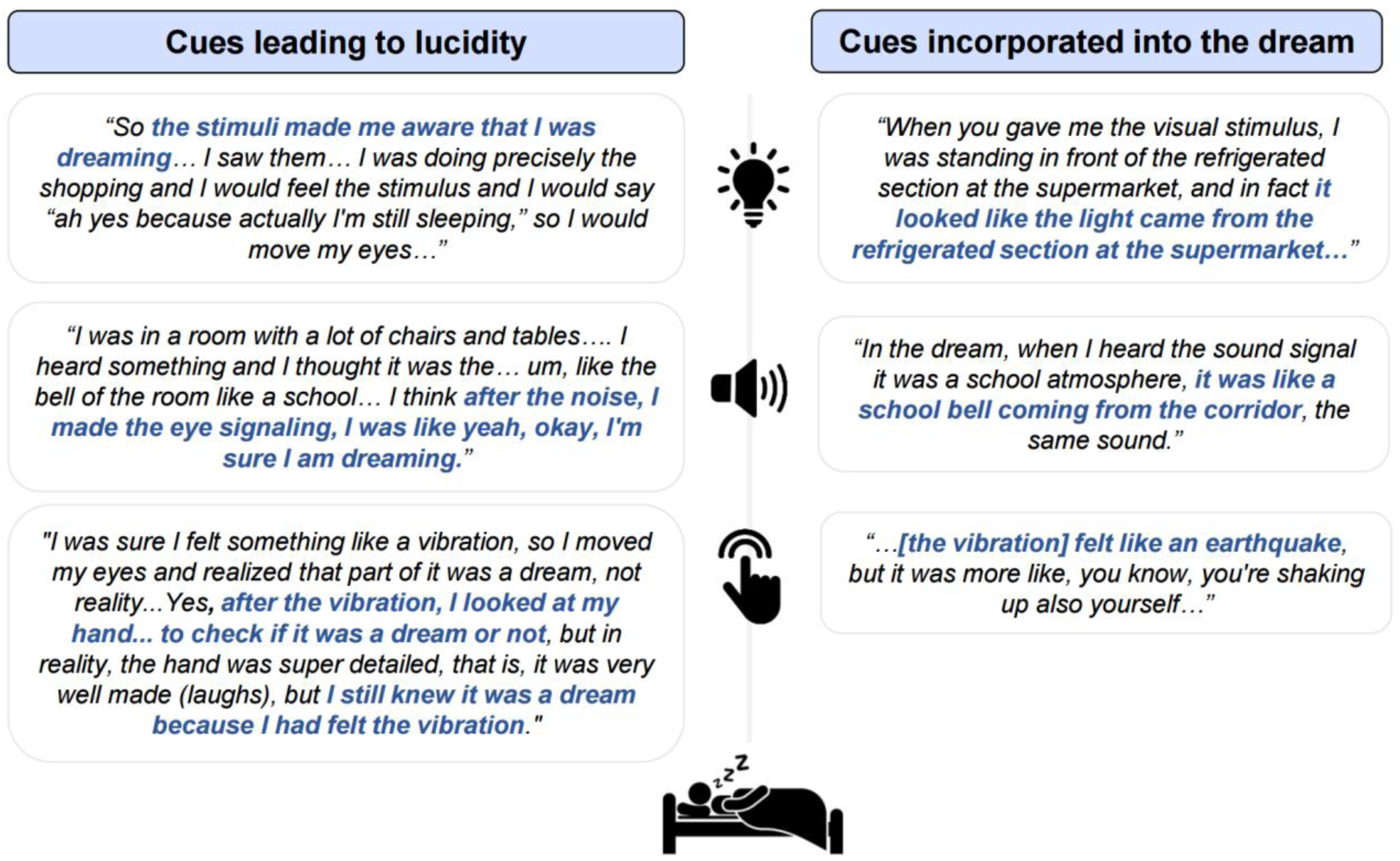
Dream excerpts showing examples of how cues presented in REM sleep led either to lucidity or were incorporated into the dream. Examples are provided for visual, auditory and tactile cues.

As self-reported by participants, 9 SVLDs (39.1% of SVLDs in stim naps) were directly initiated by perceiving cues from within the dream (5 V, 2 A, 2 T; see examples in Figure 6). Participants also self-reported being awoken by the cues in 23 (23.7%) stim periods (5 V, 9 A, 9 T), including 8 cases in which cues were not perceived in their sleep prior to their awakening. Interestingly, in the sham naps in the absence of cueing, there were 15 instances (15.3% of 98 REM sham awakenings) in which participants reported perceiving cues while asleep. Five of these sham periods were SVLD experiences, and an additional 4 were subjective LDs.

### 3.2 Sleep measures

The effects of condition and site on sleep architecture were examined using linear mixed models (*sleep variable ∼ condition + site + (1|subject)*). REM sleep duration was shorter in the stim condition (12.7 ± 10.2 min) than in the sham condition (16.2 ± 10.4 min) (p = .040); as were sleep efficiency (stim: 53.4 ± 13.6%; sham: 57.6 ± 12.5%; p = .025) and sleep maintenance efficiency (stim: 64.4 ± 14.0%; sham: 68.9 ± 12.5%; p = .021). All other sleep variables did not significantly differ by condition (all p > .162). Several sleep metrics, but not those related to REM sleep, significantly differed by laboratory site (see Supplementary Table S3).

### 3.3 Self-reported post-study effects

To assess whether the study had any carryover effects on participants’ dreams, they were asked to complete an optional follow-up survey two weeks after completing the study. Out of 49 participants who filled out the survey, 16 (32.7%) reported having had at least one lucid dream at home after the study (1 to 3 lucid dreams; average: 1.71 ± 0.80). A total of 23 (46.9%) participants reported that the study had some impact on their dream life, including higher dream recall and more vivid dreams, feeling closer to lucidity or doing reality checks within dreams, being more aware of recurring dream themes, reflecting on how dreams connect with waking life, and experiencing false awakenings. Finally, 5 (10.2%) participants reported that the study had some impact on their waking life, including doing SSILD or reality checks at home, noticing similarities between sleepiness and the LD state, and feeling more positive and focused on the present moment.

## 4. Discussion

We carried out a pre-registered multi-center study using a wearable EEG headband, dream engineering toolbox, and a combined cognitive-sensory stimulation protocol to induce lucid dreams. We tested the efficiency of pre-sleep SSILD training combined with REM sleep TLR in participants with varying prior LD experience, using both objective and subjective measures of lucidity. We recruited 60 participants across three laboratories, representing the largest sample size to date for an in-lab LD induction study.

A relatively high rate of LD occurred over the two morning naps: 73.3% of participants had at least one subjective LD, which were objectively verified with eye signals in 51.7% of participants. However, the incidence of subjective LDs did not differ between conditions (50% stim, 55% sham), nor did the incidence of SVLDs (38.3% stim, 28.3% sham). Nonetheless, SVLDs in the stim condition were almost twice as long (6.1 ± 6.9 min) compared to the sham condition (3.2 ± 5.4 min), as measured from the first to last eye signal. Accordingly, SVLDs in the stim condition included more eye signals compared to sham SVLDs, and levels of dream control were marginally higher in the stim condition than in the sham condition. Overall, our findings indicate no clear advantage of TLR over the SSILD method alone on the incidence of LDs, although it may help with prolonging LD duration and enhancing control.

Sensory stimulation in REM sleep has previously yielded varying results, from ineffective (Paul et al., 2014), to enhancing subjective LD experiences but not SVLDs (Erlacher et al., 2020) to Carr et al.’s initial TLR study, which reported a high objective success rate (50% SVLDs vs 38.3% in this study) when combining visual and auditory stimuli with pre-sleep training (Carr et al., 2023). A few key differences with the TLR protocol used in Carr et al. (2023) may explain these varying results, including the present study’s use of a within-subject design, the integration of a 30-minute SSILD training before sleep as compared to a shorter lucidity training, and the additional use of tactile cues beyond visual and auditory cues. Moreover, the Carr et al. (2023) study recruited a sample of participants with frequent nightmares, which could partly explain their higher induction rates, as nightmare distress was found to increase the likelihood of SVLDs. Nevertheless, our combined SSILD-TLR protocol (stim condition) induced relatively long lucid dreams, with an average duration of 6.1 min (first to last LRLR), compared to 1.9 min for Carr et al. (2023).

The impact of sensory cues on lucid dreaming warrants further discussion. Cues were perceived by sleeping participants in 71.1% of cued REM periods, with perception varying from incorporation into the dream narrative to recognition as external stimuli with or without an ongoing dream. Participants attributed cues as leading to lucidity in 9/23 cases, or 39.1% of SVLDs in stim naps. While REM cueing may not consistently help with lucidity initiation, it may serve as a reliable lucidity “reminder” and help sustaining lucidity for longer, with one participant reporting “*I think I continuously knew [that I was dreaming] because there are these sensory cues.*” This lucidity-reminding mechanism may also allow for a more precise and objective estimation of lucidity duration than relying solely on subjective reports.

Nonetheless, participants reported being awoken by the cues in 23 cases (23.7%). Of note, even the lowest settings allowed by the ZMax headband for vibration and light stimulation were judged too strong for some participants; this may have affected the success rate of TLR cueing compared with earlier studies. In fact, some participants described the sensory stimulation as detrimental for lucid dreaming: “*I decided to fly a lot. […] They [the stimuli] were almost annoying, […] every time a stimulus arrived, it seemed as if I had to learn to fly again. As if they were bringing me back to reality”*. Additionally, we presented sensory cues starting 20 seconds into REM periods, which may have been before REM sleep was stabilized. Standard sleep measures showed that the cueing procedure resulted in shorter REM sleep duration and lower sleep efficiency compared to naps with no cueing.

Interestingly, there were 15 cases where participants reported perceiving cues during the sham condition, including within 9 LDs, 5 of which were SVLDs. It is possible that pre-sleep exposure to cues and the expectation of cueing during sleep promoted lucidity within subsequent dreams. One novelty of this study was the extension of the TLR approach by incorporating tactile cues on the forehead alongside auditory and visual stimulation. Although the three cue types were perceived at comparable rates, the eye signalling response time was faster following tactile compared to visual cues. While a previous LD induction study found low success rates using tactile cues, their effectiveness was dependent on stimulation site (Paul et al., 2014) warranting further investigation on the role of body location.

A review (Tan & Fan, 2023) of LD induction techniques found that SSILD was a “potentially successful” method, with one home study showing a ∼17% rate of LDs across one week based on subjective reports (Adventure-Heart, 2020). It is thus noteworthy that SSILD alone in this study was associated with high rates of LDs (55% subjective LDs, 28.3% SVLDs) in the laboratory. However, in the absence of a no-intervention control condition, we cannot determine the extent to which these rates exceed baseline LD frequency; as a reference, in Carr et al. (2023) the control group had a baseline 17% SVLD rate. Likewise, we cannot determine whether high lucidity rates under SSILD alone, whether driven by the technique itself or by an expectation effect from sleeping in the lab, approached a biological ceiling that limited the detection of additional TLR benefits. Nevertheless, the observed LD incidence suggests that SSILD may be a promising induction technique.

Finally, LDs in this study were induced in participants with a wide range of prior LD experience. Although participants who had at least one SVLD in the laboratory tended to have higher prior LD experience, it is notable that 10 participants with little prior experience (<1 LD yearly) achieved an SVLD in the laboratory. This suggests that our approach may facilitate lucid dreaming across varying levels of LD experience, rather than being limited to experienced lucid dreamers.

Of note, some accounts have raised concerns regarding possible adverse impacts of lucid dreaming (Ableidinger & Holzinger, 2023; Soffer-Dudek, 2019). In this study, we included a brief follow-up assessment two weeks after participation to evaluate participants’ perceived impact of the study on their dreams and waking lives. At follow-up, 32.7% of participants indicated having at least one LD at home following the study, and 46.9% indicated some changes in their dream life, including higher recall, more vivid dreams, being closer to lucidity, doing reality checks within dreams and, in one case, experiencing a false awakening. No participants reported negative impacts attributable to the study, and overall feedback was generally positive.

Nevertheless, future research could further examine the longer-term effects of LD induction techniques, including both potential benefits and adverse outcomes.

### 4.2 Limitations

While the present study builds on previous work using a pre-registered within-subject procedure in a relatively large sample, several limitations can be acknowledged. First, the study had no passive control group, as all conditions included at least one lucidity induction technique. Thus, our current design did not allow for an assessment of the individual impact of SSILD training, or of REM cueing alone, on inducing LD. Testing these LD induction techniques in more naturalistic settings will also be crucial for assessing the impact of other factors inherent to in-lab experiments, such as sleeping in a novel environment or simply participating in a LD study.

Second, the use of EEG wearables for sleep recordings and sleep stimulation also presents both limitations and advantages. The lack of occipital EEG in ZMax may have restricted the ability to detect fluctuations in alpha bandpower, complicating the distinction between N1 and Wake, as well as the detection of arousals. Wearable devices may also limit experimental control over the intensity of sensory stimulation, increasing the likelihood of unintended arousals. Future studies would benefit from employing lower-intensity sensory stimulation to minimize sleep disruption, as well as from further investigating trait differences in sensory sensitivity (Schredl et al., 2022). Nonetheless, having wearable EEG as a minimal sensing system opens the possibility to replicate the current study over a longitudinal period at home.

### 4.3 Conclusions and future directions

In summary, LDs were successfully reported and objectively validated in the laboratory following the implementation of a combined cognitive-sensory induction protocol using wearable EEG and dream engineering tools, including among individuals with little prior LD experience. While TLR in this study did not increase LD induction rate over SSILD alone, it shows potential for enhancing the duration of lucidity. Continued development of reliable, wearable, at-home induction protocols may facilitate the use of LD for therapeutic applications and expand its utility as a scalable tool for sleep and consciousness research.

## Data Availability Statement

The data underlying this article will be shared on reasonable request to the corresponding author.

## Author Contributions

Conceptualization: MJE, LS, CPD, ED, SFS, MC, MD; Data curation: CPD, MJE, LS, TM, TB, BP, CC; Funding acquisition: MD, MC, GB, SFS; Methodology: MJE, LS, CPD, TB, ED, SFS, MC, MD; Software: MJE; Validation: CPD, MJE, LS, TM; Formal analysis: CPD, MJE, LS, TM, TB, VL; Investigation: CPD, MJE, LS, TM, VL; Supervision: SFS, MC, MD, GB; Visualization: CPD, MJE, LS, TM, E.P.; Writing - original draft: CPD, MJE, LS, TM, VL; Writing - review & editing: CPD, TM, MJE, LS, TB, ED, E.P., VL, BP, GB, PZ, NA, SFS, MC, MD.

## Acknowledgments

The authors thank Karen Konkoly, Amir Hossein Daraie, Yon Keuren and Frederik D. Weber for their valuable input in the study’s early planning stages; Marieke McKenna for recording the English audio track for lucidity training (NL site), Bianca Pedreschi for recording the Italian audio track (IT site), and Helena Deland for recording the French and English audio track (CA site).

## Study Funding

This work was supported by the PPP Allowance made available by Health∼Holland, Top Sector Life Sciences & Health (LSHM19062) (M.D.), the Natural Sciences and Engineering Research

Council of Canada (No. RGPIN-2024-04094); (M.C.) and (No. BP - 577789 - 2023); (C.P.D.) and Swiss National Science Foundation (No. P2ZHP1_195248) (SFS). G.B. is supported by an European Research Council Starting Grant (No. 948891) and by the Italian Ministry of University and Research (MUR) through the FARE 2020 programme (No. R2049C8YN3).

## Supplementary Figures

**Figure S1.**
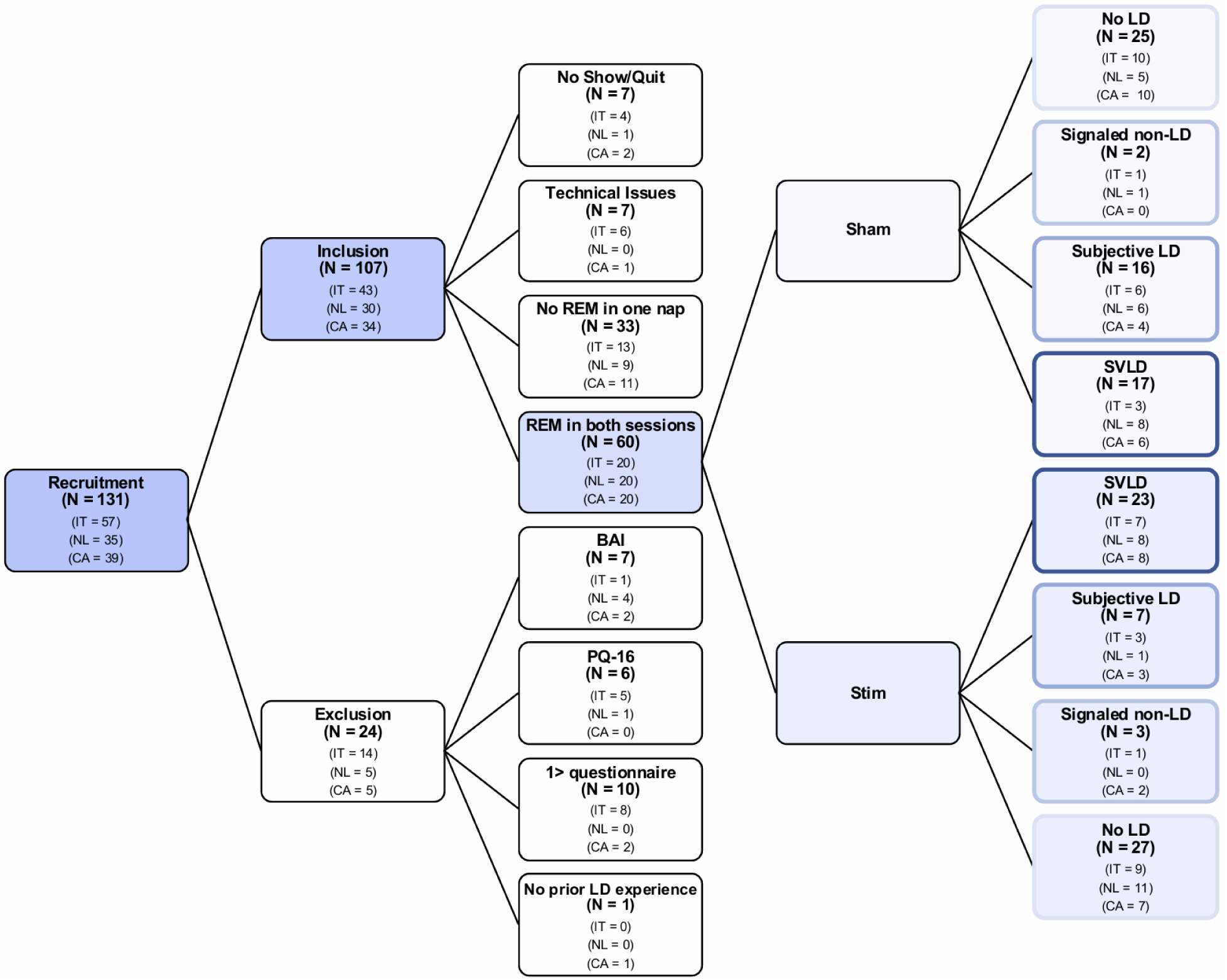
Participant attrition flowchart. Numbers are shown for the whole sample, and for the three lab sites separately.

**Figure S2.**
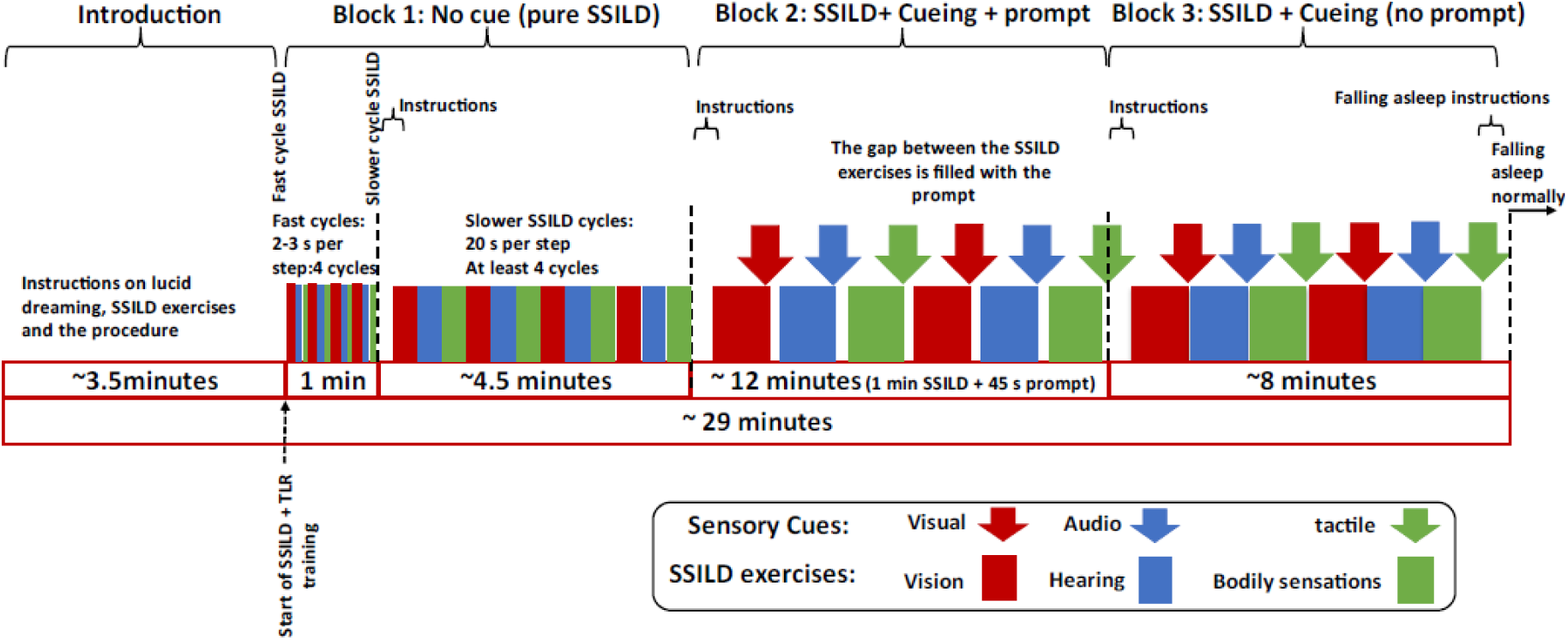
Structure and timing of the cognitive training protocol. Block 1: SSILD exercises without stimulation (fast and slower cycles). Block 2: A combination of the SSILD exercises with associated multimodal sensory stimulation (i.e., visual, audio, tactile) with verbal prompts. Block 3: A combination of the SSILD exercises with associated and multimodal sensory stimulation (i.e., visual, audio, tactile) without verbal prompts. N.B: while overall length was highly similar for all centers, the timings above correspond to the English recording, and slight block timing variations were possible due to language differences.

**Figure S3.**
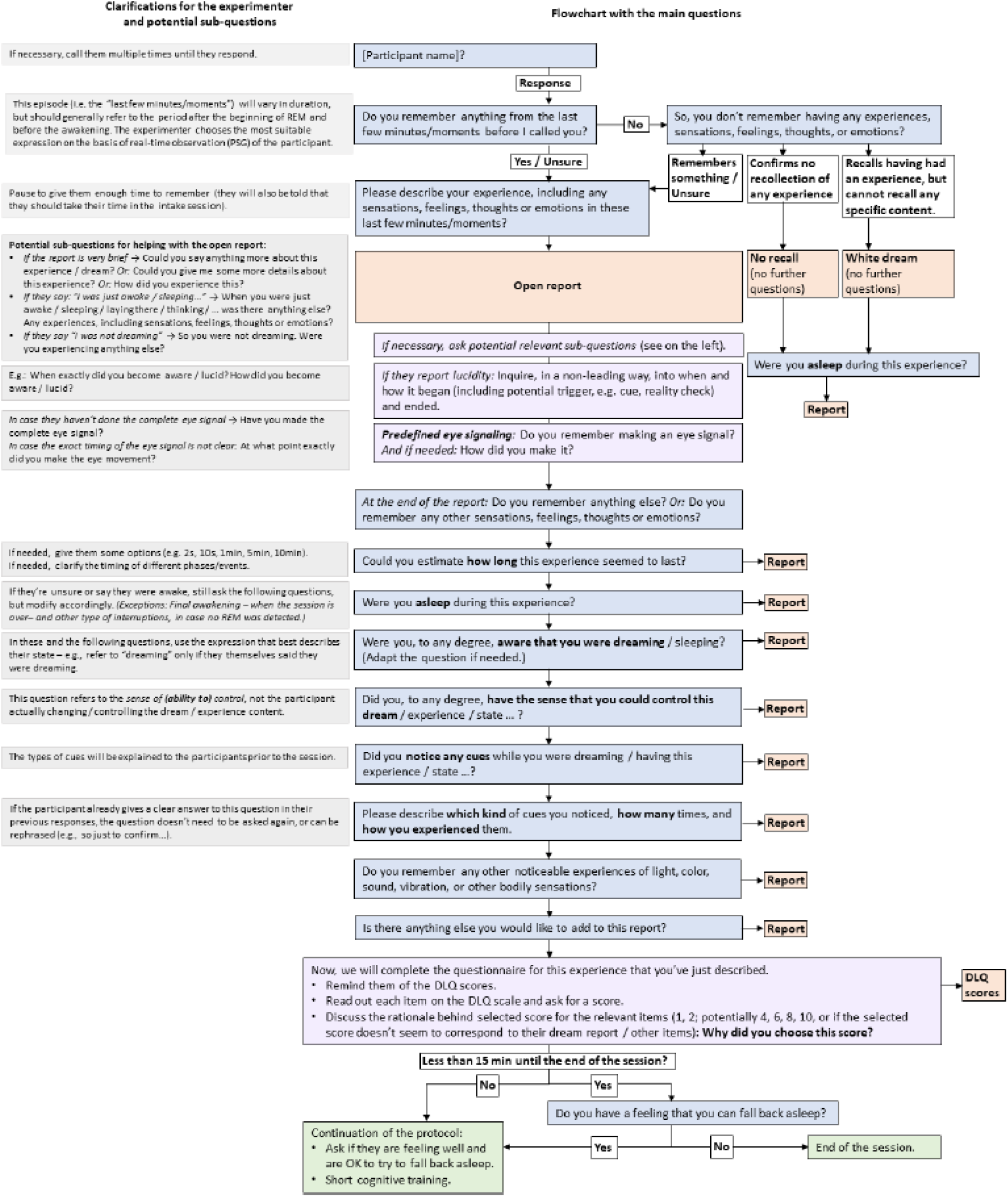
Flowchart of the semi-structured dream report interview following each experimental awakening.

**Figure S4.**
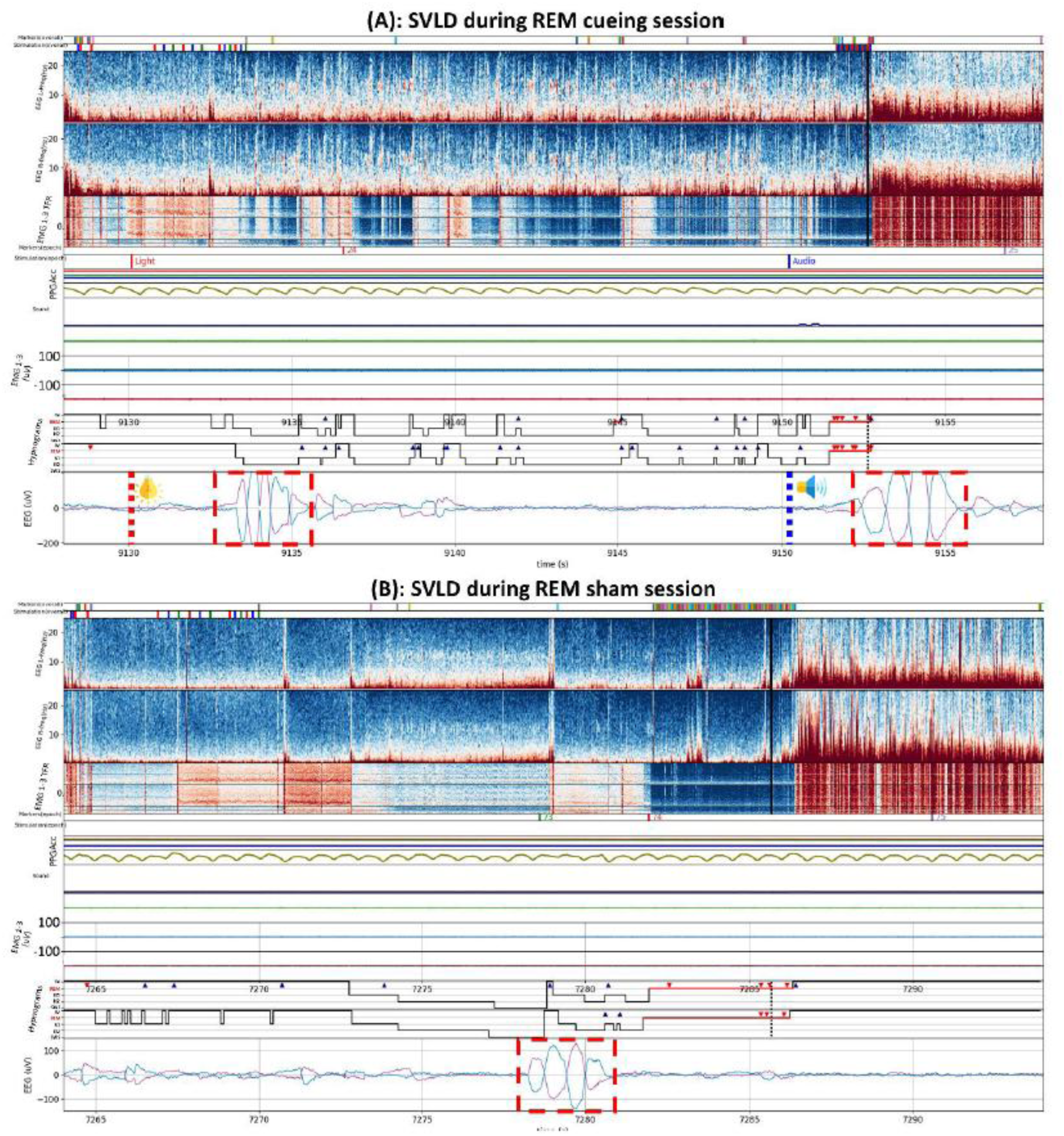
Sample SVLD cases during (A) REM cueing and (B) REM sham sessions from Dreamento user interface. An accurate interpretation of each subplot can be found in the Dreamento publication (Esfahani, Daraie et al., 2023). (A) Two subsequent predefined eye signals after the presentation of visual and auditory cues. (B) Cue-unrelated predefined eye signal. In both cases, two independent raters scored the epoch as REM sleep and agreed on scoring the eye signal as a predefined LRLR lucidity signal. The hypnograms feature dark triangles indicating arousal markers and red triangles denoting the detection of predefined eye signals. The vertical dashed line on the hypnogram represents where the current epoch of data is located with respect to the scoring. The predefined eye signals were marked using red rectangles in the EEG signal. The example is provided from the NL site.

## Supplementary Tables

**Table S1.** Cueing protocol during the initial SSILD cognitive training period. The timing of each cue within the SSILD training and their corresponding characteristics (i.e., repetition, onset/offset, intensity) are described in this table. Cognitive training blocks correspond to the ones highlighted in Figure S2. The first six cues belong to the second block (i.e., SSILD + sensory cueing + prompts), and the last six correspond to the third block of cognitive training (i.e., SSILD + sensory cueing). The cues were played in cyclic order, mirroring the SSILD exercises, and their intensities (light and audio cues) or number of repetitions (vibration cue) followed a sequentially decreasing pattern from the first to the last cycle. Individual thresholds were accounted for (see Supplementary Material 2), and stimulus intensity (light and audio cues) was adjusted based on the previously defined subjective thresholds.

| Cue number | Type | Features | Cognitive training block |
| --- | --- | --- | --- |
| 1 | Light | RED – 3 reps - 500 ms on/off - Minimum subjective intensity + 15% | 2 |
| 2 | Audio | 500-700-900 Hz , 200 ms on/off - Minimum subjective volume + 15 dBA | 2 |
| 3 | Vibration | All colors set to zero, 3 reps, 200 ms on / 500 ms off | 2 |
| 4 | Light | RED – 3 reps - 500 ms on/off - Minimum subjective intensity + 10% | 2 |
| 5 | Audio | 500-700-900 Hz , 200 ms on/off - Minimum subjective volume + 10 dBA | 2 |
| 6 | Vibration | All colors set to zero, 2 reps, 200 ms on / 500 ms off | 2 |
| 7 | Light | RED – 3 reps - 500 ms on/off - Minimum subjective intensity + 5% | 3 |
| 8 | Audio | 500-700-900 Hz , 200 ms on/off - Minimum subjective volume + 5 dBA | 3 |
| 9 | Vibration | All colors set to zero, 2 reps, 200 ms on / 500 ms off | 3 |
| 10 | Light | RED – 3 reps - 500 ms on/off - Minimum subjective intensity | 3 |
| 11 | Audio | 500-700-900 Hz , 200 ms on/off - Minimum subjective volume | 3 |
| 12 | Vibration | All colors set to zero, 1 rep, 200 ms on / 500 ms off | 3 |

**Table S2.** Cueing protocol during the short SSILD cognitive training following each awakening.

| Cue number | Type | Features |
| --- | --- | --- |
| 1 | Light | RED – 3 reps – 500 ms on/off -Minimum subjective intensity + 5 % |
| 2 | Audio | 500-700-900 Hz , 200 ms on/off - Minimum subjective volume + 5 dBA |
| 3 | Vibration | All colors set to zero, 2 reps, 200 ms on 500 ms off |

**Table S3.** Sleep metrics for the stim and sham naps.

| Sleep metric | Stim<br>(mean ± SD) | Sham<br>(mean ± SD) | Test - condition | | Test - laboratory site | | |
| --- | --- | --- | --- | --- | --- | --- | --- |
| | | | $\chi^2(1)$ | p | $\chi^2(2)$ | p | post-hoc |
| TIB (min) | 192.55 ± 16.07 | 188.69 ± 18.53 | 1.81 | .179 | 3.70 | .157 | — |
| SPT (min) | 159.08 ± 22.01 | 157.93 ± 24.87 | 0.09 | .768 | 1.61 | .447 | — |
| WASO (min) | 55.78 ± 22.49 | 49.37 ± 21.88 | 3.85 | .050 | 8.18 | .017 | IT < NL * |
| TST (min) | 103.16 ± 28.86 | 108.44 ± 25.24 | 1.74 | .187 | 5.83 | .054 | — |
| N1 (min) | 31.47 ± 19.24 | 27.98 ± 13.71 | 1.49 | .223 | 1.15 | .563 | — |
| N2 (min) | 50.73 ± 19.38 | 54.36 ± 18.61 | 1.96 | .162 | 3.50 | .174 | — |
| N3 (min) | 8.27 ± 12.13 | 9.91 ± 13.35 | 0.63 | .427 | 8.82 | .012 | CA > NL * |
| REM (min) | 12.69 ± 10.24 | 16.20 ± 10.42 | 4.21 | .040 * | 1.35 | .510 | — |
| NREM (min) | 90.47 ± 27.13 | 92.24 ± 20.73 | 0.21 | .647 | 6.09 | .048 | IT > NL * |
| SOL (min) | 19.28 ± 11.97 | 19.70 ± 14.32 | 0.05 | .822 | 30.17 | <.001 | IT > CA ***, IT > NL ** |
| N1 latency (min) | 19.48 ± 11.76 | 20.46 ± 15.88 | 0.21 | .644 | 29.89 | <.001 | IT > CA ***, IT > NL ** |
| N2 latency (min) | 38.51 ± 19.85 | 34.53 ± 15.47 | 1.97 | .161 | 11.30 | .004 | IT > CA ** |
| N3 latency (min) | 95.29 ± 48.94 | 112.07 ± 53.93 | 0.12 | .727 | 6.76 | .034 | — |
| REM latency (min) | 88.74 ± 34.28 | 83.40 ± 34.31 | 0.72 | .396 | 1.06 | .590 | — |
| N1 (%) | 31.09 ± 17.39 | 26.81 ± 13.79 | 2.89 | .089 | 0.16 | .925 | — |
| N2 (%) | 49.26 ± 14.11 | 50.35 ± 14.57 | 0.23 | .634 | 1.94 | .379 | — |
| N3 (%) | 7.51 ± 11.80 | 8.53 ± 11.54 | 0.31 | .575 | 9.36 | .009 | CA > NL * |
| REM (%) | 12.13 ± 9.08 | 14.32 ± 8.02 | 2.21 | .137 | 1.23 | .541 | — |
| NREM (%) | 87.87 ± 9.08 | 85.68 ± 8.02 | 2.21 | .137 | 1.23 | .541 | — |
| SE (%) | 53.41 ± 13.60 | 57.58 ± 12.48 | 5.01 | .025 * | 3.34 | .188 | — |
| SME (%) | 64.43 ± 14.02 | 68.86 ± 12.52 | 5.35 | .021 * | 8.02 | .018 | IT > NL * |

Sleep metrics are calculated with YASA Python Toolbox. TIB: Time in bed, SPT: sleep period time, WASO: wake after sleep onset, TST: total sleep time, SOL: sleep onset latency, SE: sleep efficiency, SME: Sleep maintenance efficiency. Measures of Latency SWS was calculated in a subsample of 21 participants who reached SWS in both naps. Values are represented in the form of mean ± standard deviation. Linear mixed models: sleep variable ∼ condition + site + (1|subject). Fixed effects were tested using Type II Wald chi-square tests. Post-hoc comparisons using Tukey correction. \*: $p < .05$ , \*\*: $p < .01$ , \*\*\*: $p < .001$ .

## Supplementary Methods

### Signal acquisition and sleep scoring

#### PSG systems

Electromyographic (EMG) channels in the IT and NL sites were recorded using site-specific software and systems (NL: BrainVision and BrainAmp ExG, Brainproducts GmbH, Gilching, Germany; IT: g.Recorder for g.tec Suite and g.USBamp Research, g.tec medical engineering GmbH, Graz, Austria). At the CA site, a standard polysomnography montage was applied using Natus SleepWorks and Natus Embla® NDx Amplifier (Middleton, USA).

#### Calibration

A calibration period started with one minute of eyes-closed resting wakefulness to establish baseline physiological measures. To synchronize EMG with headband recordings in offline Dreamento, participants were asked to clench their teeth three times in a row at the beginning and end of the recording, as well as after any awakening or interruption. Participants were also asked to perform the LRLR signal with their eyes closed as a reference to recognize their LRLR signals during sleep.

#### Sleep scoring

The sleep scorers were blind to the condition and real-time annotations. Inter-rater agreements on ZMax scoring (naps from NL and IT sites) were computed using Cohen’s kappa statistics. Due to occasional difficulty distinguishing N1 from Wake in the absence of ZMax occipital EEG channels–particularly when frontal alpha activity was not prominent–these stages were merged for inter-rater agreement analyses. When agreement fell below 80%, scorers re-evaluated conflicting epochs until consensus was reached. Kappa scores for the NL site reached 0.82 ± 0.12 (0.69 ± 0.12 without combining W and N1 stages). Kappa scores for the IT site reached 0.83 ± 0.11 (0.67 ± 0.13 without combining W and N1 stages).

### Qualitative analyses of dream reports

Dreams were scored to assess the occurrence of dream experiences often associated with LD, according to the following definitions:

- Lab incorporation dreams (LID): participants dreamt about the laboratory, such as being in or around the laboratory, perceiving electrodes and other lab objects, or interacting with experimenters.^58^
- False awakenings (FA): participants dreamt of waking up within their dreams.^59^
- Sleep misperception (SM): cases where participants reported being awake despite physiological signals indicating a sleeping state.^60^
- Sleep paralysis (SP): the experience of being temporarily unable to move or talk (muscle atonia), often occurring in the transitions between sleep and wakefulness and sometimes accompanied by vivid sensory or mental experiences.^61^
- Out-of-body experiences (OBE): dreams where the participants observed their sleeping body from a third-person perspective.^62^

### Evaluation of individual stimulation intensity level

Before sleep, the minimum subjective thresholds for visual stimuli (with eyes closed), auditory stimuli (against low-level white noise), and background white noise were individually adjusted based on the subjective appraisal of the participant, as follows:

a. Background white noise was set to an unobtrusive volume level, i.e., 35 dbA. The subject was asked whether the volume level was endurable; otherwise, the volume was lowered by 5 dBA. Background white noise was initiated at the start of the experimental nap procedure, before the pre-sleep cognitive training, and was kept constant until the end of the nap session.
b. The audio cue volume was initially set to 40 dBA. While participants were in closed-eye resting wakefulness, the experimenter invited them to judge whether this volume was perceptible and estimate whether it could be arousing if presented during sleep. If the participant agreed that this volume level was non-arousing, the minimum audio volume was set to 40 dBA; otherwise, the experimenter lowered the minimum audio cue volume by 5 dBA and repeated the process until the participant agreed upon the selected volume. The resulting volume level was set as ‘minimum subjective audio volume’.
c. The lowest light cue intensity was set at 1% (500 ms on/off) in the Dreamento software. Participants were asked to indicate whether they could perceive the light during closed-eye resting wakefulness. If the light cue was not perceived, the experimenters increased the light intensity by 5% until the subjective lowest threshold was reached. The resulting intensity of light was set as ‘minimum subjective light intensity’.
d. The lowest vibration intensity was set at 1 repetition (200 ms on / 500 ms off) in the Dreamento software. Perception of the vibration was verified for all participants during closed-eye resting wakefulness.

## Supplementary Material: Full lucidity training script (long and short versions)

### Long version

The cognitive training was presented through a pre-recorded audio track. Each center used different languages for the recording (NL: English; IT: Italian; CA: French and English) while keeping the content, length, and structure identical. The audio track starts with a general definition of lucid dreaming, provides a summary of the cognitive training protocol, and then describes the different fast (2-3 seconds of focus on each of the senses in cyclic order, i.e., vision, hearing, and bodily-sensation exercises) and slow (20 seconds of focus on each senses) cycles of SSILD training in detail (Supplementary Figure S2, Block 1). The first training block instructs participants to perform 4 uncued fast cycles, followed by 4 uncued slower cycles.

Afterward, the audio tracks describe the association of the stimuli with the SSILD cycles: light cues with the vision exercises, audio cues with the hearing exercises, and vibration cues with bodily-sensation exercises. Each SSILD exercise (vision, hearing, and bodily-sensations, respectively) lasts one minute and concludes with the administration of the corresponding sensory cue by the experimenter to remind the participant to keep a lucid mindset. The cues also mark the transition from one exercise to the next, with a light cue indicating the end of the vision exercise and the start of the hearing exercise, an audio cue signaling the completion of the hearing exercise and the initiation of the bodily-sensation exercise, and a tactile cue marking the conclusion of the cycle and the start of a new one. During this block, each sensory cue is accompanied by a verbal prompt describing the targeted lucid mindset, as in Carr et al. (2020). Finally, the last block of the cognitive training consists of a repetition of the previous cued SSILD section, with the sole exception that no verbal prompts are played after each cue. As a consequence, participants are expected to transition autonomously from one exercise to another whenever they perceive a cue. Participants are also reminded to perform the predefined LRLR eye movement signaling in cases of lucidity or stimulus perception after falling asleep.

[Bloc 1 - Introduction]

*A lucid dream is a dream in which you are aware of the fact that you are dreaming while you are still asleep. With this realization, you can sometimes influence or control what happens in a dream. In this experiment, you’ll get instructions on how to practice becoming lucid while awake*.

*For lucid dreams to occur, you need to train your mind and body into a subtle state that is optimized for lucid dreaming. This involves focusing on your vision, hearing, and bodily sensations in cycles. From this point, we guide you through the training process and describe the practicing cycles. The cycles always start with a vision exercise, then continue towards the hearing exercise, and finally end with the bodily-sensation practices. We describe each of the vision, hearing, and bodily-sensation exercises now*.

[Vision exercise]

*For the vision exercise, you should keep your eyes closed and focus all your attention on the darkness behind your closed eyelids. Keep your eyes completely still and totally relaxed. You might see colored dots, complex patterns, images, or maybe nothing at all. It doesn’t matter what you can or cannot see – just pay attention in a passive and relaxed manner and don’t try to see anything*.

[Hearing exercise]

*For the hearing exercise, we want you to shift all of your attention to your ears. You might be able to hear the faint sounds of traffic or the wind from outside. You might also be able to hear sounds from within you, such as your own heartbeat or a faint ringing in your ears. It doesn’t matter what, if anything, you can hear – just focus all of your attention on your hearing*.

[Bodily sensations exercise]

*For the bodily sensations exercise, you should shift all of your attention to sensations from your body. Feel the weight of your blanket, your heartbeat, the temperature of the air, etc. You might also notice some unusual sensations such as tingling, heaviness, lightness, spinning sensations, and so on. If this happens, simply relax, observe them passively and try not to get excited*.

*Before starting with the first exercise, I would like to mention that you may have intrusive thoughts during this process. For instance, you may think of what you need to buy or do after this experiment. It doesn’t matter if the intrusive thoughts come to your mind, but, what matters is that you can intentionally let them go and focus on your mind and body*.

[Bloc 2 - SSILD]

**1 minute of fast cycles practice followed by 4 minutes of short cycles practice.**

[4 fast cycles]

“*Now you should start with the first step of the training. Practice **four fast cycles** during which you spend **only 2-3 seconds** focusing each on the vision, hearing, and bodily sensations. You don’t have to count the seconds, but you should complete at least four cycles during this time. You can start now*.

[1 min of silence]

[4-6 slow cycles]

*You can stop now. At this point, you should perform **four to six slower** cycles that approximately take **20 seconds** focusing each on the vision, hearing, and bodily sensation steps. Again, you don’t have to count the seconds, but you should complete at least four cycles during this time. You can start now*.

[4 min of silence]

**[Bloc 3 - SSILD + TLR + prompts]**

*“Now, I want to train your mind to recognize the flashing lights, beeping sounds, and vibration cues as <u>lucidity cues</u> so that when one is played during your sleep, you will become lucid in a dream. While you rest here, we are going to play the cues at intervals. Whenever you hear, see, or feel one of the cues, you should remain in the same position with your eyes closed, but you will become lucid by attending to where your mind has been, attending to your body, and attending to your surroundings. The same as before, each cycle starts with vision training. You should continue the vision practice until you see a light cue. Then a prompt will be played to help you focus. Once the prompt is ended you should move forward to the hearing exercise. You should continue the hearing practice until you hear an audio cue. Again, a prompt will be played to help you focus. Once the prompt is ended you should move forward to the bodily-sensations exercise. The vibration cue will indicate when the exercise is ended and then you should start a new cycle. Please note that, as a response to the cues, you do not need to do the Left-right-left-right eye signals while training, only do the eye signals when you become lucid in a dream*.

*You can start with vision practice, now… ”*

[2 cycles, i.e. 6 x cues (Light-audio-tactile-light-audio-tactile) – 1 min intervals]]

[Prompt]

*“As you notice the cue, you become lucid. Bring your attention to your thoughts … <u>[pause]</u> …, notice where your mind has wandered …<u>[pause]</u>… Now observe your body … <u>[pause]</u> …, your sensations … <u>[pause]</u> …, and your feelings… <u>[pause]</u> … Observe your breathing…<u>[pause]</u>… remain critically aware, lucid, and notice how aspects of this experience are in any way different from your normal waking experience. <u>[pause]</u>”*

[The next exercise starts by mentioning the type of the exercise, i.e., vision, hearing, or bodily-sensation]

*“You can start with vision/hearing/bodily-sensations practice, now…”*

**[Bloc 4 - SSILD + TLR (no prompt)]**

The last block of the cognitive training employed an identical SSILD procedure coupled with sensory cueing, with the sole exception that no verbal prompts were played to guide the participant. As a consequence, the participant was expected to transition independently to the subsequent exercise following each cue. The participant was also reminded of doing the predefined LRLR eye movement in case of lucidity during sleep

[Instructions after the last prompt (6th cue)]

“*The cues will continue to be played in intervals over the next 6 minutes. The prompts, however, will not be played anymore. You should continue to practice becoming lucid by focusing on your vision, hearing, and bodily sensations, the same as before and again in cycles. We keep sending you the cues when you need to move from one exercise to another. Pay attention to your mind, your body, and your surroundings. Notice how aspects of your experience are in any way different from your normal waking experience. At the end of this block, when the cues are stopped, you can fall asleep normally and you don’t have to do the exercises anymore. Please keep in mind that when the cognitive training is ended, you should do the left-right-left-right eye signaling in three cases: (1) when you become lucid in a dream, (2) as a response to the cues while being lucid, and (3) approximately every 30 seconds while you are lucid, even though you do not perceive any cue*.

*Now you can start with the vision exercise…”*

[2 cycles, i.e., 6 cues (2 from each type) are played with 1-min intervals]

### Short version

[4 fast cycles]

*We will do a very short round of training, again. The same as before, we want you to practice **four fast cycles** during which you spend only **2-3 seconds** focusing each on the vision, hearing, and bodily sensations. You don’t have to count the seconds, but you should complete at least four fast cycles during this time. You can start now*.

[1 minute of silence] [1 slow cycle]

*At this point, you should perform **one slower cycle** of approximately **20 seconds** focusing on each on the vision, hearing, and bodily sensation steps. Again, you don’t have to count the seconds, but you should complete at least one cycle during this time. You can start now*.

[1 minute of silence]

[3 slow cycles with cues]

*Now, the same as before, we will play the cues at intervals. You should continue to practice becoming lucid by focusing on your vision, hearing, and bodily sensations in cycles. Whenever you need to move from one exercise to the next, you will receive the corresponding cue. At the end of this block, when the cues are stopped, you can fall asleep normally and you don’t have to do the exercises anymore. Please keep in mind that when the cognitive training is ended, you should do the left-right-left-right eye signaling in three cases: (1) when you become lucid in a dream, (2) as a response to the cues while being lucid, and (3) approximately every 30 seconds while you are lucid, even though you do not perceive any cue*.

*You can start with vison practice now*.

[1 cycle : 3 cues]

## Notes

### Competing Interest Statement

The authors have declared no competing interest.

### Summary of Updates

Results from third site of this multi-center study were added.

